# Protein classification using modified *n*-*gram* and *skip*-*gram* models

**DOI:** 10.1101/170407

**Authors:** S M Ashiqul Islam, Benjamin J Heil, Christopher Michel Kearney, Erich J Baker

## Abstract

**Motivation:** Classification by supervised machine learning greatly facilitates the annotation of protein characteristics from their primary sequence. However, the feature generation step in this process requires detailed knowledge of attributes used to classify the proteins. Lack of this knowledge risks the selection of irrelevant features, resulting in a faulty model. In this study, we introduce a means of automating the work-intensive feature generation step via a Natural Language Processing (NLP)-dependent model, using a modified combination of N-Gram and Skip-Gram models (m-NGSG).

**Results:** A meta-comparison of cross validation accuracy with twelve training datasets from nine different published studies demonstrates a consistent increase in accuracy of m-NGSG when compared to contemporary classification and feature generation models. We expect this model to accelerate the classification of proteins from primary sequence data and increase the accessibility of protein prediction to a broader range of scientists.

**Availability:** m-NGSG is freely available at Bitbucket: https://bitbucket.org/smislam/mngsg/src

**Supplements:** link to supplementary documents

## 1 INTRODUCTION

It is well appreciated that primary polypeptide sequence informs higher order protein structure. The primary sequence provides the blueprint which encodes the purpose of the protein, ultimately determining the proteins characteristics, functions, subcellular localization and interactions (Pour-El and American Chemical Society, 1979). However, classical approaches using primary sequence alignment for the prediction of remote homology detection are problematic due to low signal to noise ratios in polypeptide strings (Teichert *et al.*, 2010). To circumvent this problem, non-alignment based methodologies are being investigated to demonstrate remote homology (Bonham-Carter *et al.*, 2014; Vinga and Almeida, 2003; Liu *et al.*, 2014; Du *et al.*, 2014). Here we illustrate a novel approach that relies on Natural Language Processing (NLP) to produce generalized feature sets for machine learning classification of protein characteristics.

A polypeptide string can be treated as a text string where hidden information is deciphered by implementing NLP techniques. Generating *n*-*grams* (Cavnar *et al.*, 1994) and *skip*-*grams* (Guthrie *et al.*, 2006) from text documents is a feature extraction method which can produce meaningful information for machine learning (ML) classification algorithms (Cavnar *et al.*, 1994; Guthrie *et al.*, 2006), and has been used for the categorization and sorting of documents based on their subject matter (Tan *et al.*, 2002; Hu and Liu, 2004; Pang *et al.*, 2002). Treating a primary protein sequence as a textual string is a natural extension of this approach. Indeed, text mining has been used previously for protein clustering and classification, protein-protein interaction (PPI), protein folding, and cnRNA identification (Zeng *et al.*, 2015). Linguistic methodologies based on primary sequence features have also been applied in areas of secondary structure prediction (Ding *et al.*, 2014b).

Sequence classification using supervised and unsupervised machine learning methods is becoming popular due to algorithm accessibility in conjunction with increasing amounts of available biological data. Recent work in this area includes the classification of protein structure (Islam *et al.*, 2015), localization (Yu and Hwang, 2008), function (Cai *et al.*, 2003), family (Chou, 2005) and protein-protein interaction (PPI) (Zhao *et al.*, 2012; Yu and Hwang, 2008) based on primary sequence. These studies consistently report that ML approaches are superior to alignment based predictions when deriving protein characteristics from primary sequence, and perform effectively in protein groups with low sequence similarity. However, the success of ML models depends heavily on training data, feature extraction, classifier algorithm selection and optimization.

Among these steps, robust results are disproportionately influenced by feature selection. Thus, substantial effort is required to obtain meaningful features from protein data. While universal methods for feature extraction are problematic due to the wide range of classification strategies, several generalized feature generation methods have been proposed. Many of these methods aim to address specific classification problems (Islam *et al.*, 2015; Bock and Gough, 2001; Dyrlv Bendtsen *et al.*, 2004), while others may be implemented as semiautomated feature generators. For example, amino acid composition (Verma and Melcher, 2012) and pseudo-amino acid composition (Du *et al.*, 2014) based feature extraction schemes have been successfully used to solve a range of classification problems (Garg *et al.*, 2005; Xu *et al.*, 2013; Qiu *et al.*, 2016; Tiwari, 2016). There are also hybrid feature generation strategies which include both generalized and data specific feature selection methods (Sharma *et al.*, 2013; Chaudhary *et al.*, 2016; Ramaprasad *et al.*, 2015). In each case, however, manual intervention is required to produce the optimal set of features.

Using *n*-*grams* and *skip*-*grams* in biological applications driven by ML is not without precedent. For example, the *n-gram* model has been used to classify protein sequences into superfamilies using extreme machine learning (Cao and Xiong, 2014). Homology between proteins with low sequence similarity has also been successfully revealed using distances between *Top*-*n*-*gram* and amino acid residue pairs (Liu *et al.*, 2014). *Spaced words* is a derivative of *n*-*gram* feature selection in biological sequence analysis where the letters of one or more indices in each word are replaced by blanks except the first and last letters. This method of feature extraction is used along with another method called *kmacs* to perform alignment-free comparison in both DNA and protein sequences (Horwege *et al.*, 2014).

Through the application of a modified NLP *n*-*gram* and *skip*-*gram* (m-NGSG) approach, we have developed a primary protein sequence feature selection method that is fully automated, and agnostic to peptide function or chain size. A meta-comparison of logistic regression mediated classification approaches exploiting our feature generation method with other published models illustrates enhanced functional and structural binary and multi-class classification accuracy in every instance. Without the requirement of expert intervention for optimal feature selection, it is hoped that this automated approach will reduce the time needed to employ ML classification strategies for protein prediction.

## 2 METHODS

### 2.1 Feature generation, vectorization and model construction

The *n*-*gram* and *k*-*skip*-*bi*-*gram* profiles are initially extracted from each candidate protein sequence. They are given a position identity with respect to the C-terminus of the protein sequence. Thereafter, modifications of the length of *k*-*skip*-*bi*-*grams* and positional identity are performed to obtain potential motifs (or words). Finally, the motifs are vectorized to construct feature vectors with a simultaneous noise filtration. The length of initial *n*-*gram* and *k-skip-bi-gram* motifs, and amplitude of their modification are determined by six parameters (described in Supplementary Table S1.). The parameters are optimized using a modified grid search algorithm (see Algorithm 1 and 2 in Supplementary text) depending on the training set of a five-fold cross-validation using a logistic regression classifier. As the modified grid search is seeded using different initial *n*-*grams*, they are defined as *seeds* in this study (see the methods section in Supplementary Text for details).

### 2.2 Meta-comparison

The performance of m-NGSG was compared with other methodologies that use generalized or data-specific feature extraction methods for model construction. Comparison models were chosen based on the availability of benchmark data reported by those models, the diversity of protein characteristics classified, and the ability of the model to report functional or structural classification of proteins with regard to their sequence. The performance was compared with the published models using logistic regression (Table 1).

**Table 1.** Description of the models those are used in meta-comparison.

| Model Name | Dataset Description | Original Feature Extraction Method | Classifier | Ref |
| --- | --- | --- | --- | --- |
| Subchlo | Two pair of training datasets of protein sequences from 4 classes based on their localization in chloplast. One dataset includes the raw sequences of proteins here annotated as Subchlo raw. Another dataset consists of sequences less than 60% identity annotated as Subchlo60 | Pseudo Amino acid composition (PseAAC) | Evidence theoretic K-nearest neighbor(ET-KNN) with a jackknife cross validation. | Du *et al.* (2009) |
| osFP | One pair of training and testing dataset of protein sequences from 2 classes (monomer vs oligomer) based on oligomeric states of proteins. We selected the benchmark protein sequences less than 95% identity. | AAC/DPC/TPC, AC, CTD, Ctriad, QSO, PseAAC | Features having a threshold 0.7 for the Pearson correlation coefficient were removed. Decision tree with ten-fold cross validation following splitting the whole training set into 80% and 20% training and test set respectively. This process was repeated for 100 time to get an unbiased confidence interval of the accuracy. | Simeon *et al.* (2016) |
| iAMP-2L | One pair of training and test dataset of two classes of protein sequences based on antimicrobial activity | PseAAC | fuzzy k nearest neighborhood (FKNN) with a Jackknife cross validation. | Xiao *et al.* (2013) |
| Cypred | One pair of training and test dataset consist of two classes of sequences based on cyclic and noncyclic structure. | AAC, cyclicpeptide specific motifs | SVM with (RBF) kernel coupled with 10-fold cross validation. | Kedariseti *et al.* (2014) |
| PredSTP | One training dataset divided into two classes based the cysteine bonding pattern in the 3D structure of the proteins. | normalized distance between cystine pairs explained in [ref] along with hydrophobic, hydrophilic, neutral and count of some other amino acids. | SVM with RBF kernel coupled with 200-fold cross validation. | Islam *et al.* (2015) |
| TumorHPD | Two training datasets annotated as TumorHPDn1 and TumorHPD 2 in this paper. The sequences are divided in two classes based on their affinity to tumor cells. TumorHPD 1 consists the raw protein sequences while TumorHPD 2 consists only the sequences not more than 10 amino acids long. | AAC, DPC, BPP | SVM with 5-fold cross validation. | Sharma *et al.* (2013) |
| HemoPI | Three pairs of training and test set annotated as HemoPI, semiHemoPI and nonHemoPI two in the paper which contains hemolytic, semihemolytic and nonhemolytic peptides, respectively. Here we compared model that classifies the raw hemolytic and nonhemolytic peptides annotated as HemoPI 1, and the model that classifies hemolytic and semihemolytic peptides annotated as HemoPI 2 | AAC, DPC, BPP | SVM with 5-fold cross validation. | Chaudhary *et al.* (2016) |
| IGPred | One pair of training and test set those are divided into two classes those fall into two groups: immunoglobulin and nonimmunoglobulin | PseAAC followed by ANOVA based feature Selection technique | SVM with RBF kernel coupled with a Jackknife cross validation. | Tang *et al.* (2016) |
| PVPred | One pair of training and test set those are divided into two classes those fall into two groups: virion and nonvirion . | g-gap dipeptide composition plus PseAAC followed by ANOVA based feature Selection technique. | SVM with RBF kernel coupled with a Jackknife cross validation. | Ding *et al.* (2014a) |

Abbreviations: PseAAC = Pseudo amino acid composition; APC = Amino acid composition; DPC = Dipeptide composition; TPC= Tripeptide composition; AC = Auto correlation; CTD = Composition, Transition, Distribution; Ctriad= Conjoint triad; QSO=Quasi-sequence-order; BPP = binary profile pattern (presence or absence of a motif of interest)

In addition, m-NGSG was evaluated with a linear kernel SVM classifier on the Subchlo raw and Subchlo60 datasets (Du *et al.*, 2009) to demonstrate that the m-NGSG feature extraction algorithm works equally well with other classifiers.

The m-NGSG feature extraction method was also compared with other generalized extraction methods. Here, the Quantitative Structure-Property Relationship (QSAR) based feature generation method (Simeon *et al.*, 2016) was implemented on the Subchlo60 data set from Subcholo model, and compared with the m-NGSG feature generation model. A logistic regression classifier was used with a regularization parameter of C=1, and evaluated using Jackknife, 5-fold, and 10-fold cross validation.

## 3 RESULTS

### 3.1 Parameter optimization analysis

This study illustrates that *n*-*gram* and *skip*-*gram* text mining approaches can be exploited to develop a generalized feature extraction method for protein classification. *N*-*gram* and *skipgram* models are not used directly; rather, the models are modified according to six parameters based on sequence (Supplementary Table S1.). The parameters themselves are optimized by using the modified grid search-based algorithm m-NGSG (see Algorithm 2 in Supplementary text) and compared to 12 benchmark datasets. In each case, the automated generalized feature extraction algorithm obtained features that outperformed the originally published feature sets for linear regression.

For the benchmark datasets iAMP-2L, Cypred, TumorHPD 1, TumorHPD 2, IGPred and PVPred, the optimization strategy for m-NGSG reported the same parameters (see Supplementary Fig.S1.) with identical accuracy (Supplementary Fig.S2) regardless of the initial seed, indicating convergence in these data sets. For the subchlo raw training set, parameters n, k, and y showed variation with some seeds, (see Supplementary Fig.S1). Overall, the subchlo raw training set accuracy for different seeds ranged from 89%-89.70% (Supplementary Fig.S2). For the subchloro60 training set, parameters *n*, *k*, *y*, and *c* demonstrated variability over the first four seeds and then became stable while the accuracy ranged from 65.76% to 68.07%. In the PredSTP training set, there was slight variation in parameter *n*, *k* and *y* which was also reflected in the variation of accuracies for the corresponding seeds. Parameters for the HemoPI 1 training set varied for seed three, and training set HemoPI 2, which classifies between hemolytic and semihemolytic peptides, presented variation in parameters *n*, *k*, *kp*, *y* and *c* for seed 3, 4 and 5 (see Supplementary Fig. S1 and Fig.S2).

The goal of parameter optimization is to identify parameters that contribute to the best accuracy after five-fold cross validation. Although the principle approach is a modified grid-search, it demonstrates an ability to converge on accuracy regardless of initiating seeds. Supplementary Fig.S3 illustrates the convergence characteristic of the optimization algorithm which calculates the mode value of accuracies generated from different seeds against the percent change of the accuracies from each seed for a specific training set when compared to the mode accuracy. Flat areas in supplementary Fig.S3 indicate low percentage change compared to the mode which suggests convergence.

### 3.2 Meta-comparison of prediction performance on benchmark datasets

Once the parameters were optimized for each benchmark training set, the reported accuracy was compared to the m-NGSG model built with the optimized feature set. A logistic regression classifier was used for all models. To compare the cross-validation accuracy, we mimicked the approach published as part of the original dataset, either five-fold, ten-fold or jackknife validation.

#### 3.2.1 Subchlo

Subchlo is a multi-class classifier designed to predict the localization of chloroplast proteins. Subchlo raw is a dataset of protein sequences based on their location in chloroplast and the Subchloro60 dataset represents proteins with approximately 60% sequence identity. Subchlo raw and Subchlo60 were both cross-validated by a jackknife method in the original publication, resulting in a combined accuracy of 89.69% and 67.18%, respectively. The accuracy of the m-NGSG model is 91.59% and 73.92% for the same datasets (see Supplementary Table S2). This indicates a 2.12% and 7.73% increase of accuracy by our model compared to the reported model for the two given datasets (Figure 1A).

**Fig. 1.**
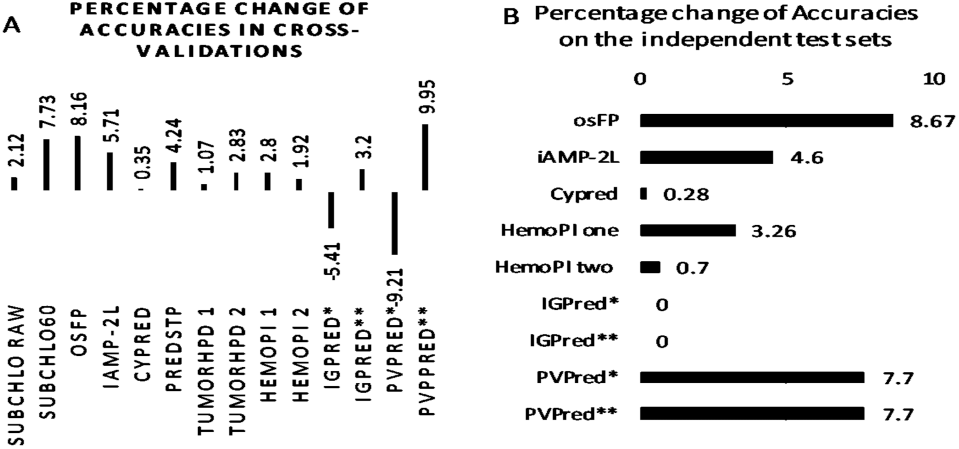
The percentage changes of accuracies m-NGSG in crossvalidation compared to the original models for each dataset. IGPred* and PVPred* shows the comparative accuracy changes without feature selection while IGPred** and PVPred** shows accuracy changes after mimicking the feature selection method of the original model (A). The percentage changes of accuracies m-NGSG on the independent test sets (depending on availability) compared to the original models. IGPred * and PVPred* shows the comparative accuracy changes without feature selection while IGPred** and PVPred** shows accuracy changes after mimicking the feature selection method of the original model (B).

#### 3.2.2 osFP

The osFP model classifies fluorescent proteins into monomer or oligomeric states. In the original study, different QSAR (Quantitative StructureActivity Relationship)-based feature selection models were investigated. The best model yielded an average of 72.13% and 72.89% accuracy for the training and test sets after 100 iterations (see Suplementary Table S5). In contrast, m-NGSG generated an average of 78.02% and 79.21% accuracy for the same sets, yielding an 8.16% and 8.6% increase of accuracy respectively (Figure 1). To confirm the superiority of m-NGSG model over the QSAR based feature selection method, we also performed a comparison on Subchlo60 dataset. The comparison demonstrated that m-NGSG’s performance is better than that of other feature generation methods (see Supplementary Table S6 and Supplementary Fig.S4).

#### 3.2.3 iAMP-2L

The iAMP-2L model classifies antimicrobial peptides from nonantimicrobial peptides. Supplementary Table S2 and Table S3 illustrates the increased performance of m-NGSG over the iAMP-2L when using jackknife cross validation method. The accuracy of m-NGSG on the training set was 91.25%, yielding a 5.71% rise over the previously reported accuracy. When we used m-NGSG to evaluate the performance on the benchmark independent test set, we achieved a 4.6% rise from the accuracy reported by the original model (Figure 1A).

#### 3.2.4 Cypred and PredSTP

Both Cypred and PredSTP classify proteins based on their structural characteristics. While Cypred performed comparably to m-NGSG (99.20% accuracy after 10-fold cross-validation in the original publication vs 99.53% for m-NGSG), m-NGSG did provide a modest 0.35% increase. On a bechmark out of sample test data set, the m-NGSG model narrowly outperformed Cypred by 0.28%. On the other hand, a comparison on training set cross validation accuracy between PredSTP and m-NGSG produces a 2.50% gain of accuracy from the original model (see Supplementary Table S2 and S3, and Figure 1).

#### 3.2.5 TumorHPD 1 and 2

TumorHPD classifies tumor homing peptides to identify analogs of tumor homing ability. Two training sets were used to generate the models: raw tumor homing peptides, *TumorHPD 1*, and tumor homing peptides less than or equal to ten residues long *TumorHPD 2*. Among three different generation methods they used (Sharma *et al.*, 2013), amino acid composition yielded the best accuracy 82.52% and 80.28% for the training set TumorHPD 1 and 2, respectively. The accuracy of m-NGSG the same datasets were 83.40% and 82.55%, respectively (see Supplementary Table S2) which using logistic regression yielding a 1.07% and 2.83% rise from the original model(Figure 1A).

#### 3.2.6 HemoPI 1 and 2

HemoPI 1 model classifies hemolytic and nonhemolytic proteins, while HemoPI 2 classifies hemolytic and semi hemolytic peptides. The performance data for the training and test sets were available for the models developed from hybrid feature sets. The original model searched for the best accuracy by considering whole proteins and fractions of the proteins. Here, we compared the m-NGSG accuracy with only the whole length proteins. Our model generated 97.97% accuracy for HemoPI 1 and 79.5% accuracy (Supplementary Table S2) for HemoPI 2 training sets offering a 2.8% and 1.92% increase from the original models respectively. When we compared m-NGSG on the benchmark independent test sets, it achieved an increase of 3.26% and 0.7% for HemoPI 1 and HemoPI 2 respectively (Figure 1).

#### 3.2.7 IGPred and PVPred

IGPred predicts immunoglobulin protein, and PVPred predicts virion proteins from primary sequence data. The size of these these proteins is very different from that of previously classified proteins. Immunoglobulin and virion proteins have very long sequences. In both models important features were selected using ANOVA analysis before performing the jackknife cross validation. Therefore, we also performed jackknife cross validation with and without an ANOVA-based feature selection method where we used the minimum number of features offering the best cross-validation accuracy (see Supplementary Fig.S5). The accuracy of m-NGSG model was 100% with ANOVA-based feature selection, and 92.60% with jackknife cross validation (Supplementary Table S2), while the accuracy of the original IGPred model with jackknife test was 96.60%. The accuracy for the independent test set was 100% regardless the model (Supplementary Table S3). For PVPred, the accuracy of jackknife cross validation with and without feature selection was 89.25% and 77.19% respectively, with corresponding accuracies of 90% and 93.33% on the benchmark independent test sets. The original feature selection assisted model showed 85.02% accuracy for jackknife cross validation and 86.66% accuracy for the independent test set (Supplementary Table S3)

## 4 DISCUSSION

The crucial steps of machine learning-based classifications are the selection of datasets that unambiguously represent informative classes, creation of meaningful features from the dataset that can optimally correlate to different classes, and an appropriate choice of machine learning algorithms which effectively classify the data based on the data points and descriptors. Predicting protein characteristics from primary sequence is becoming popular as appropriate data sources experience rapid growth and computer libraries for machine learning algorithms become accessible to bench biologists. However, generating effective features from protein sequences continues to require enormous manual intervention, and automated approaches have narrowly scoped structure prediction. Chemical property-based feature generation algorithms and dipeptide or tripeptide motif-specific approaches (Chaudhary *et al.*, 2016; Kedarisetti *et al.*, 2014) account for the the majority of these feature generation methods. In particular, Pseudo Amino Acid Composition (PseAAC) has been the most frequently used approach to classify proteins per their functional properties(Xiao *et al.*, 2013; Mohabatkar *et al.*, 2013), subfamilies(Chou, 2005), interactions with other proteins(Jia *et al.*, 2015) and subcellular localizations(Lin *et al.*, 2008). Methods that classify based on physicochemical or biochemical properties rely heavily on the AAindex database (Kawashima and Kanehisa, 2000).

However, as protein sequences are strings of amino acid residues, they can be treated as normal text that can be interpreted through by NLP-based techniques. The m-NGSG algorithm presented herein generates features in a text mining manner where words are artificially generated from protein sequences using modified *n*-*gram* and *skip*-*gram* models. The models themselves are optimized based on the combination of six parameters (Supplementary Table S1.). NLP processing of protein strings creates a corpus of words that is subsequently used for vectorization to generate features for each individual data point. To fully automate the classification process, a modified grid search algorithm is employed to obtain the optimal values of the six parameters. The parameter optimization itself is performed after 5-fold cross validation to confirm the whole training set is not exposed to the classifier during the optimization step, limiting the risk of bias during the meta-comparison. Moreover, all the optimization was done with a logistic regression classifier with the same regularization parameter value to avoid disparity in this step.

Interestingly, although the optimization algorithm primarily depends on a modified grid search, in most cases parameters converge to a single value regardless of the initial seed (Supplementary Fig.S1). Also, in many cases, the different starting seeds yield the same accuracy (Supplementary Fig.S2). These outcomes indicate that the optimization algorithm searches for the maximum value while retaining the ability to converge.

A collection of contemporary models were chosen for meta-comparison based on their diversity of classification topic (such as functional, structural and subcellular localization), database size, sequence length and feature selection methods (Table 1). Benchmark training datasets from comparison model publications were used (Supplement datasets). With the exclusion of the osFP dataset, the meta-analysis comprised six of the eleven independent test sets (five were unavailable). In the case of osFP, the original dataset was divided into training and test sets and ten-fold cross validation was performed only on the training set. For the models without an independent test set, evaluation with cross validations on the benchmark datasets were performed as an adequate replacement to reveal the comparative performance between the models.

The m-NGSG model outperformed cross-validation accuracy of each model it was compared against, with the increase in accuracy ranging from 0.35%-9.95% over the original models (Figure 1A). Moreover, we observed up to an 8.67% increase in accuracy over the original model when compared to independent test sets (Figure 1B). As shown in Figure 1A, the cross validation accuracy of IGPred and PVPred without feature selection was considerably less than the original model where ANOVA based feature selection was performed before the execution of jackknife cross validation accuracy, while the same ANOVA based feature selection method in m-NGSG model displayed higher jackknife cross validation accuracy on the same training set. The accuracy on the independent test set demonstrated a 0% and 7.7% increment from the original IGPred and PVPred, respectively, regardless of which feature selection was used (Figure 1B). This result illustrates that feature selection method followed by cross validation test biases the cross validation process without improving the performance of a model.

Two of the models in our meta-comparison (TumorHPD 1 and HemoPI 1) reported accuracy based on protein fragments as well as whole protein sequence. While fragment-based models provided for slightly better accuracy, they are not included in this study because they are beyond the scope of demonstrating a generalized feature extraction method on whole sequences.

The Subchlo60 and osFP datasets were used to compare the performance of the m-NGSG model with motif composition, represented by AAC/DPC/TPC, and chemical property-based feature generation methods, represented by AC, CTD, Ctriad, SOCN, QSO and PseAAC methods (Supplementary Table S5 and S6). The m-NGSG model demonstrates a 2.12% increase over the PseAAC-based model on the Subchlo raw dataset. However, with the low sequence identity Subchlo60 data set we observed a 7.73% increase in accuracy (Figure 1A). This result indicates that m-NGSG performs comparatively better than chemical property-based method when the sequence identity in the training dataset is lower. In addition, the accuracy of m-NGSG outperformed all of the competitors in the osFP model (Supplementary Table S5), illustrating the robustness of the m-NGSG model for feature generation when compared to presently available approaches.

During the vectorization step, instead of counting feature frequencies for each data-point, only the binary profile of the features were considered. This approach reduces the complication of the model, subsequently minimizing the chance of over-fitting.

To maintain the equality the in comparison at the cross validation step with the training set, we adopted the same cross validation method with the same dataset reported in the original method. Logistic regression with the default values from scikit-learn library was used as classifier for both the optimization and meta-comparison step. We used the same classifier in the meta-comparison process because the ultimate goal of the study is to elucidate the potential effectiveness of the m-NGSG feature generation method, not the classification algorithm. Even though the classifier selection is beyond the scope of the study, we reported the accuracy of cross validation with SVM for Subchlo raw and Subchlo60 dataset which shows a better performance than logistic regression for both datasets (Supplementary Table S4). This data indicates that the m-NGSG feature generation method is compatible with multiple classifiers.

## 5 CONCLUSION

The meta-comparison results outlined in this study illustrate that the m-NGSG is an effective fully automated feature generation method. This model will benefit the machine learning-based protein classification community, particularity those interested in classification based on primary protein sequence. It is expected that m-NGSG will significantly reduce the work load at the feature generation step regardless of protein characteristics and sequence size. Moreover, by analyzing the feature importance, the distinguishing part of the sequence (motif) in a protein class can be revealed, which is often difficult to discover using multiple sequence alignment.

## SI Method

### Feature extraction

#### Binary profile of n-grams in a protein sequence

*N*-*grams*, strings of contiguous sequences consisting of *n* items, are valuable features extracted from text or speech, and are useful in NLP and sentiment analysis (Socher R *et al.*, 2013; Ghiassi M *et al..*, 2013; Cui H, Mittal V and Datar M, 2006) Given that a primary protein sequence can be treated as a string of amino acids, *n*-*gram*-based feature extraction methods can be applied to predict functionality from a sequence. Interestingly, *n*-*grams* from a protein sequence also offer biologically meaningful information, as each *n*-*gram* represents a protein sequence motif. *N*-*gram* motifs provide information helpful in inferring protein functionality, and can be represented as:

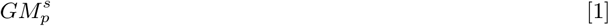

where GM stands for *Gram Motif*, and *s* is a positive integer not longer than the length, *L*, of the corresponding protein sequence (*s*|*s* ∈ ℕ, *s* <= *L*). *s* = 0 represents a null motif, *s* = 1 represents all single residue motifs (uni-grams), *s* = 2 represents all dipeptide motifs (bi-grams excluding their uni-gram components), and *s* = *n* represents all n-peptide motifs. *p* is the permutation index of the participating residue(s) parameterized by *s*. Since there are 20 different amino acids, there can be 20^*s*^ different values of *p* for an s-gram. For example, if we consider the amino acid sequence MISHW, then M is one of the 20 possible elements of uni-gram (*s* = 1) as *p* = 20^*s*^ = 20^1^ = 20. Similarly, MI is one of the 400 possible elements of dipeptides (*s* = 2) as *p* = 20^*s*^ = 20^2^ = 400.

#### Binary profile of k-skip-bi-grams

Skip-grams are a technique largely used in the field of speech processing that allow items, or in our case substrings, to be ignored during processing (Mikolov T *et al.*, 2013a; Mikolov T *et al.*, 2013b; Bian J *et al.*, 2014). In m-NGSG we adopted the *k*-*skip*-*bi*-*gram* approach where the skip distance, *k*, allows a total of *k* or fewer skips to construct the bi-gram. For example, for protein sequence MISHW, the 2-skip-bi-grams will be MI, IS, SH, HW, MXS, IXH, SXW, MXXH and IXXW where skips are represented by *X*. The *k* = 0 skips are MI, IS, SH, and HW, the *k* = 1 skips are MXS, IXH, SXW, and the *k* = 2 skips are MXXH, IXXW. This approach can be useful in comparing *k*-length mutational events across protein sequences. In order to avoid duplicating features extracted with the *n*-*gram* method, we exclude the motifs produced where *k* = 0.

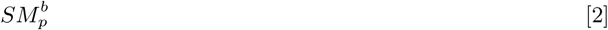

SM stands for *Skip Motif* and *b* is the number of skips between two amino acids. *b* is a positive integer that is at most two less than the length of the protein sequence (*b*|*b* ∈ ℕ, *b* ≤ *L* – 2). *b* = 0 represents no skips between a specific permutation of two residues, *b* = 1 represents one skip, and *b* = 2 represents two skips. *p* is the permutation index of the participating residue(s) parameterized by *s*. Since there are 20 different amino acids, there can be 20^2^ different values of *p* for a given value of *b*.

#### Modification of skips in k-skip-bi-gram motifs

The m-NGSG employs a modification of the *k*-*skip*-*bi*-*gram* model that allows buffering on the number of skips. That is, after obtaining the exact number of skips from a *k*-*skip*-*bi*-*gram*, an estimated number of skips is determined as:

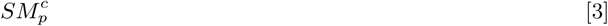

where *c* represents the estimated number of skips based on the given parameter *a*, and *b* is the number of skips in a motif as determined from the *k*-*skip*-*bi*-*gram.*

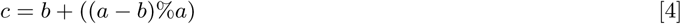

**Table S1.** Description of parameters employed in m-NGSG (modified n-gram skip-gram) based feature generation from an individual sequence.

|  |  |
| --- | --- |
| $n$ | determines the maximum length of an $n$-gram motif |
| $k$ | determines the maximum number of skips in a $k$-skip-bi-gram motif |
| $np$ | determines the maximum length of an $n$-gram motif that gets a positional value |
| $kp$ | determines the maximum skips in a $k$-skip-bi-gram motif that gets a positional value |
| $y$ | determines the positional buffering parameter in both $n$-gram and $k$-skip-bi-gram motifs |
| $c$ | determines the skip buffering parameter in $k$-skip-bi-gram motifs |

For example, if X represents a single skip, the motifs MXXH and MXH are considered unique without buffering. However, if the skips are buffered by 2 (*a* = 2), the buffered skip value of motif MXXH will be *c* = 2 + ((2 – 2)%2) = 2, yielding the original motif MXXH. On the other hand, skip buffering MXH gives the value *c* = 1 + ((2 – 1)%2) = 2, and yields a new motif MXXH. This motif is different from the original MXH, but is identical to the previous example MXXH. In this way, the buffered skip model can account for insertion/deletion events.

#### Modification of estimated C-terminus position in n-grams and k-skip-bi-grams

During feature extraction from a protein sequence m-NGSG determines the relative position of the motifs with respect to the C-terminus. *N*-*gram* or *k*-*skip*-*bi*-*gram* motifs are tagged with a maximum position identity, noted as *s*^th^ gram (for *n*-*gram*) and *b^th^*-*skip*-*bi*-*gram*(*k*-*skip*-*bi*-*gram*), respectively. This position is measured after obtaining the exact distance from the C-terminus and applying a buffering distance to capture shared positional identity for *n*-*gram* motifs,

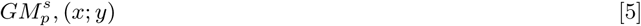

and *k*-*skip*-*bi*-*gram* motifs,

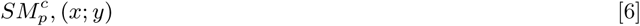

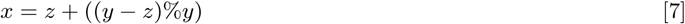

where *x* represents the distance identity of motif 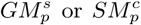 based on the given parameter *y*, and *z* is the distance of the onset of the 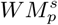 motif from the C-terminus of the sequence buffered by *y*. m-NGSG initializes *y* based on *y*_0_, defined by *ModifiedGridSearch*, and increases with the length, *l*, of the motif, as:

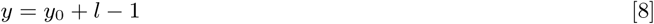

As an example, if we consider NTerm-AYHGFTVCKY-CTerm as a protein sequence, then two tyrosines will be members of the set of uni-gram motifs, and should be considered as identical. However, if we choose to account for position, each will be assigned position identity information as defined by equation (5). If the initial buffer value *y*_0_ equals 5 then the positional identity of the first Y and the last Y will be *x* = 9 + ((5 – 9)%5) = 10 and *x* = 1 + ((5 – 1)%5) = 5, respectively. Here the distance of first Y is 9 and the second Y is 1 from the C-terminus. In this way, rather being identical, the tyrosines will be recorded as Y10 and Y5 in the feature set. This approach can be generalized to *n*-*grams.* The bi-gram AY has a positional identity of 12, because its onset is 10 residues away from the C-terminus, and the buffer value will be 6 because *y*_0_ is 5 and the length of the motif is 2.

### Feature selection and model construction

Ultimately, six parameters determine the final set of features to be generated from a given sequence (Table S1). The feature extraction algorithm generates descriptors (motifs) from a list of protein sequences, which function as words in a document. To reduce noise, words that make up more than 30% of the corpus and words that appear less than 3 times are removed as an alternative to *tf*-*idf* (Joachims T 1996). Next, the model creates a sparse matrix using a vectorization method where each of the retained words or motifs composes a vector. The value of the vectors for data-points in the sparse matrix describes the presence or absence of the feature in a corresponding data-point. In other words, each row of a vector reports the presence of a selected motif in a protein sequence. Finally, a logistic regression model (Ruczinski I et al.,2003) is trained with the training data set, and its accuracy is calculated with five-fold cross-validation. The model construction scheme is done in python 2.7 using the numpy, pandas and scikit-learn packages (Pedregosa F *et al.*, 2011). When running logistic regression, a regularization constant of 1 and default parameters are used.

### Parameter optimization algorithm

The feature generation function depends on the six parameters described in Table S1. Here, a modified grid-search optimization algorithm, Algorithm 2, chooses parameters for generalized classification problems based on the accuracy of five-fold cross-validation using a logistic regression model. Briefly, it iterates over pairs of parameters to maximize accuracy, using maximal previous knowledge to inform future iterations. Each grid-search is initiated from a value of the parameter for the *n*-*gram* motifs (*n*) which is referred to here as the *seed.*

#### Algorithm 1 Logistic Regression Accuracy

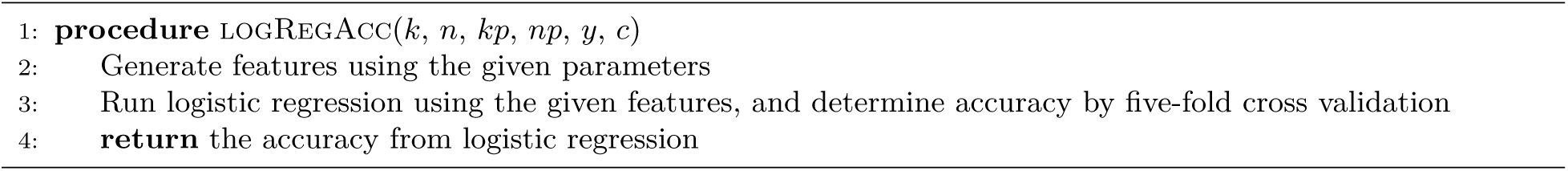

#### Algorithm 2 Modified Grid Search

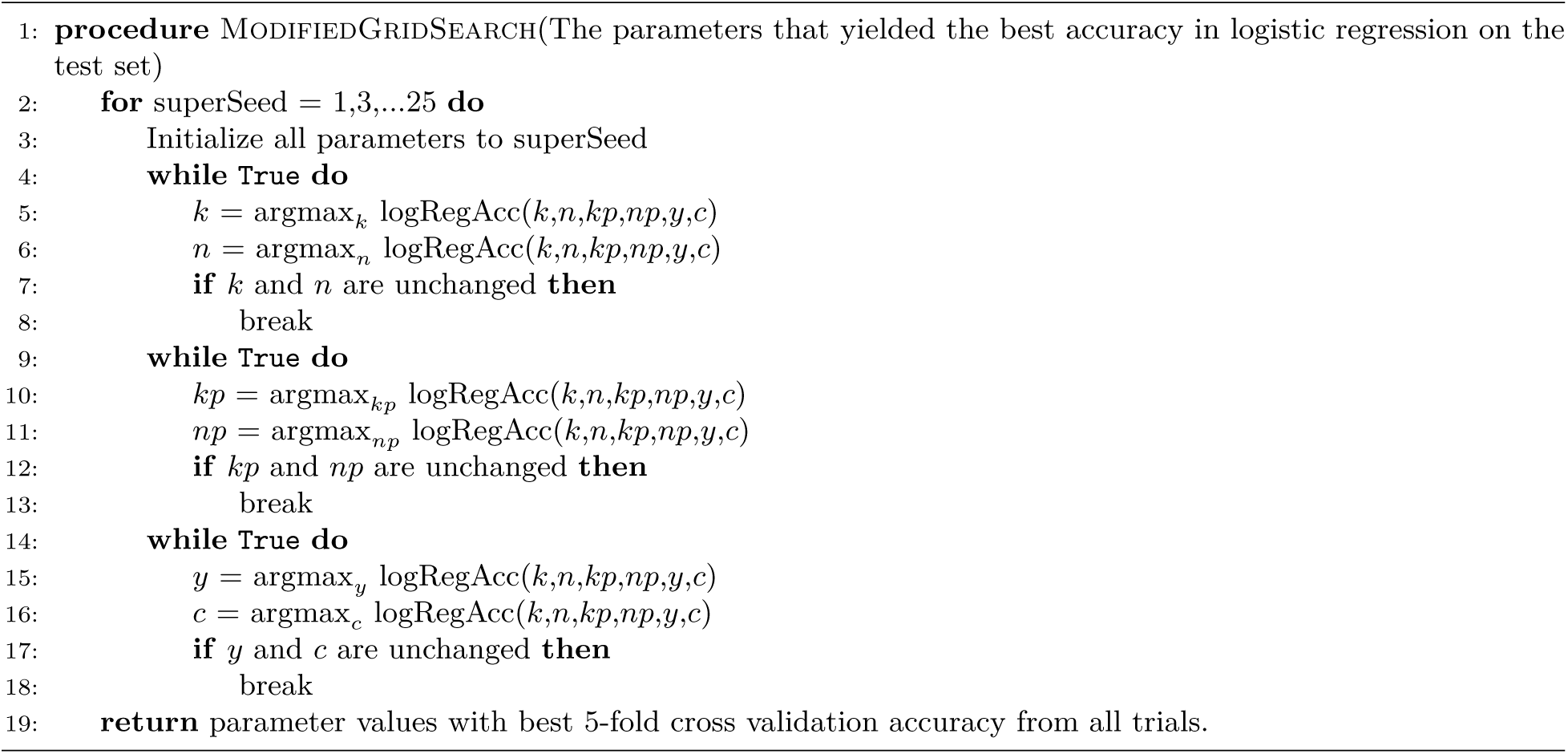

**Table S2.** Comparison between cross-validation accuracies reported on different benchmark training datasets and the corresponding accuracies achieved employing the m-NGSG model. Accuracies are displayed in percentage values.

| Classification dataset | Reported accuracy on the training set | m-NGSG accuracy on the training set |
| --- | --- | --- |
| Subchlo raw | 89.69 | 91.59 |
| Subchlo60 | 67.18 | 73.92 |
| iAMP-2L | 86.32 | 91.25 |
| Cypred | 99.2 | 99.55 |
| PredSTP | 94.3 | 96.66 |
| TumorHPD 1 | 82.52 | 83.4 |
| TumorHPD 2 | 80.28 | 82.55 |
| HemoPI 1 | 95.3 | 97.97 |
| HemoPI 2 | 78 | 79.5 |
| IGPred | 96.92 | 91.66 |
| IGPred with feature selection | 96.9 | 100 |
| PVPred | 85.02 | 77.19 |
| PVPred with feature selection | 85.02 | 93.48 |

**Table S3.** Comparison between accuracies reported on independent test sets and the corresponding accuracies achieved employing the m-NGSG model. Accuracies are displayed in percentage values.

| Classification dataset | Reported accuracy on the test set | m-NGSG accuracy on the test set |
| --- | --- | --- |
| iAMP-2L | 92.23 | 96.47 |
| CypredL | 98.7 | 98.98 |
| HemoPI 1 | 96.4 | 99.54 |
| HemoPI 2 | 75.7 | 76.23 |
| IGPred | 100 | 100 |
| IGPred with feature selection | 100 | 100 |
| PVPred | 86.66 | 93.33 |
| IGPred with feature selection | 86.66 | 93.33 |

**Table S4.** Comparison among the models on Subchlo raw and Subchlo60 datasets. LR and SVM stands for logistic regression and support vector machines classifiers, respectively. The comparison shows the accuracies for separate localization based classes along with the overall accuracies. Accuracies are displayed in percentage values.

| Locations | Subchlo raw original method | Subchlo raw m-NGSG LR | Subchlo raw m-NGSG SVM | Subchlo60 original method | Subchlo60 m-NGSG LR | Subchlo60 m-NGSG SVM |
| --- | --- | --- | --- | --- | --- | --- |
| Stroma | 78.87 | 78.87 | 78.87 | 67.35 | 69.38 | 65.30 |
| Thylakoid lumen | 55 | 63.33 | 66.66 | 43.18 | 54.76 | 59.52 |
| Thylakoid membrane | 96.12 | 98.64 | 98.06 | 83.72 | 88.97 | 88.97 |
| Envelope | 84.44 | 80.00 | 84.44 | 40 | 51.38 | 53.84 |
| Overall | 89.69 | 91.59 | 92 | 67.18 | 73.92 | 75.09 |

**Table S5.** Comparison of evaluation matrices of the m-NGSG model with the feature generation methods used in osFP dataset. Accuracies are displayed in percentage values.

| Methods | CV Accu-<br>racy% | CV MCC | Test set<br>Accu-<br>racy% | Test set<br>MCC |
| --- | --- | --- | --- | --- |
| AAC/DPC<br>/TPC | 72.13 ±<br>4.18 | 0.42 ±<br>0.08 | 72.89 ±<br>7.08 | 0.43 ±<br>0.15 |
| AC | 70.71 ±<br>4.45 | 0.38 ±<br>0.09 | 70.30 ±<br>8.55 | 0.38 ±<br>0.18 |
| CTD | 69.40 ±<br>4.95 | 0.39 ±<br>0.10 | 70.18 ±<br>7.79 | 0.38 ±<br>0.17 |
| Ctriad | 68.64 ±<br>5.99 | 0.34 ±<br>0.12 | 71.26 ±<br>8.36 | 0.40 ±<br>0.17 |
| QSO | 68.98 ±<br>4.21 | 0.34 ±<br>0.09 | 69.93 ±<br>6.90 | 0.37 ±<br>0.14 |
| PseAAC | 69.39 ±<br>4.97 | 0.35 ±<br>0.10 | 69.67 ±<br>8.03 | 0.36 ±<br>0.17 |
| **m-NGSG** | 78.02 ±<br>0.93 | 0.50 ±<br>0.02 | 79.21 ±<br>1.47 | 0.54 ±<br>0.03 |

**Table S6.** Comparison of accuracies on the Subchlo60 dataset obtained by different generalized feature generation methods and m-NGSG. Comparisons were showed for Jackknife, 5-fold and 10-fold cross-validations. Logistic regression was used as classifier for all models. Accuracies are displayed in percentage values.

| Method | Accuracy % |  |  |
| --- | --- | --- | --- |
|  | Jackknife | 5-fold CV | 10-fold CV |
| AAC/DPC/TPC | 49.41 | 45.00 | 49.61 |
| AC | 59.14 | 58.07 | 59.61 |
| CTD | 58.36 | 57.3 | 55.38 |
| Ctriad | 52.52 | 50.38 | 52.69 |
| SOCN | 57.19 | 55.76 | 57.69 |
| QSO | 49.41 | 45.00 | 49.61 |
| m-NGSG | 73.92 | 70.00 | 72.69 |

**Fig. S1.**
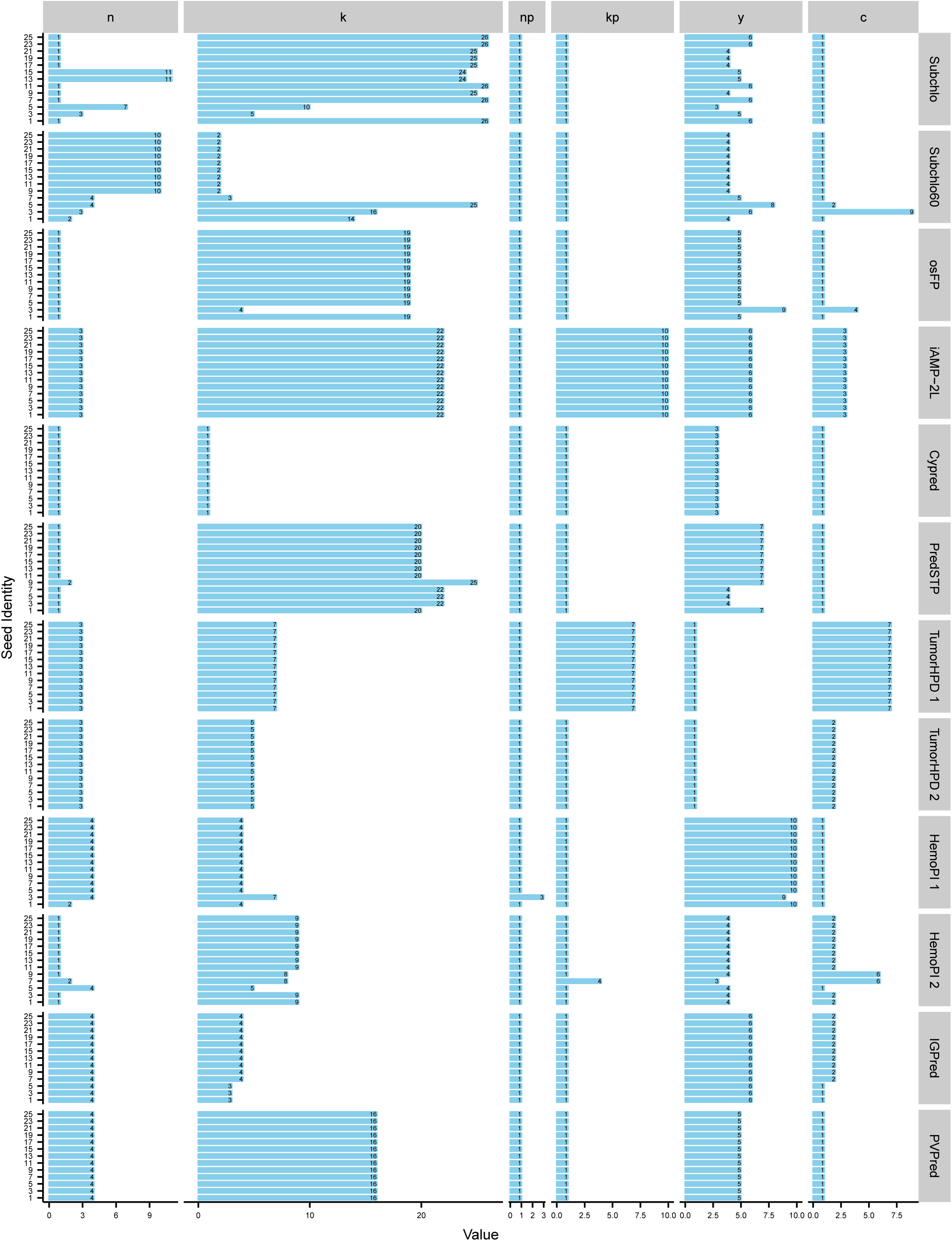
Illustrates the optimum values of individual parameters generated from each seed for a specific dataset. The values of the six parameters were optimized from thirteen different seeds (the initial value of n). Each column in the panels (n, k, np, kp, y, c) assign the parameters and each row represent individual dataset. The X axis shows the value of the parameters, while the y axis represents an individual seed.

**Fig. S2.**
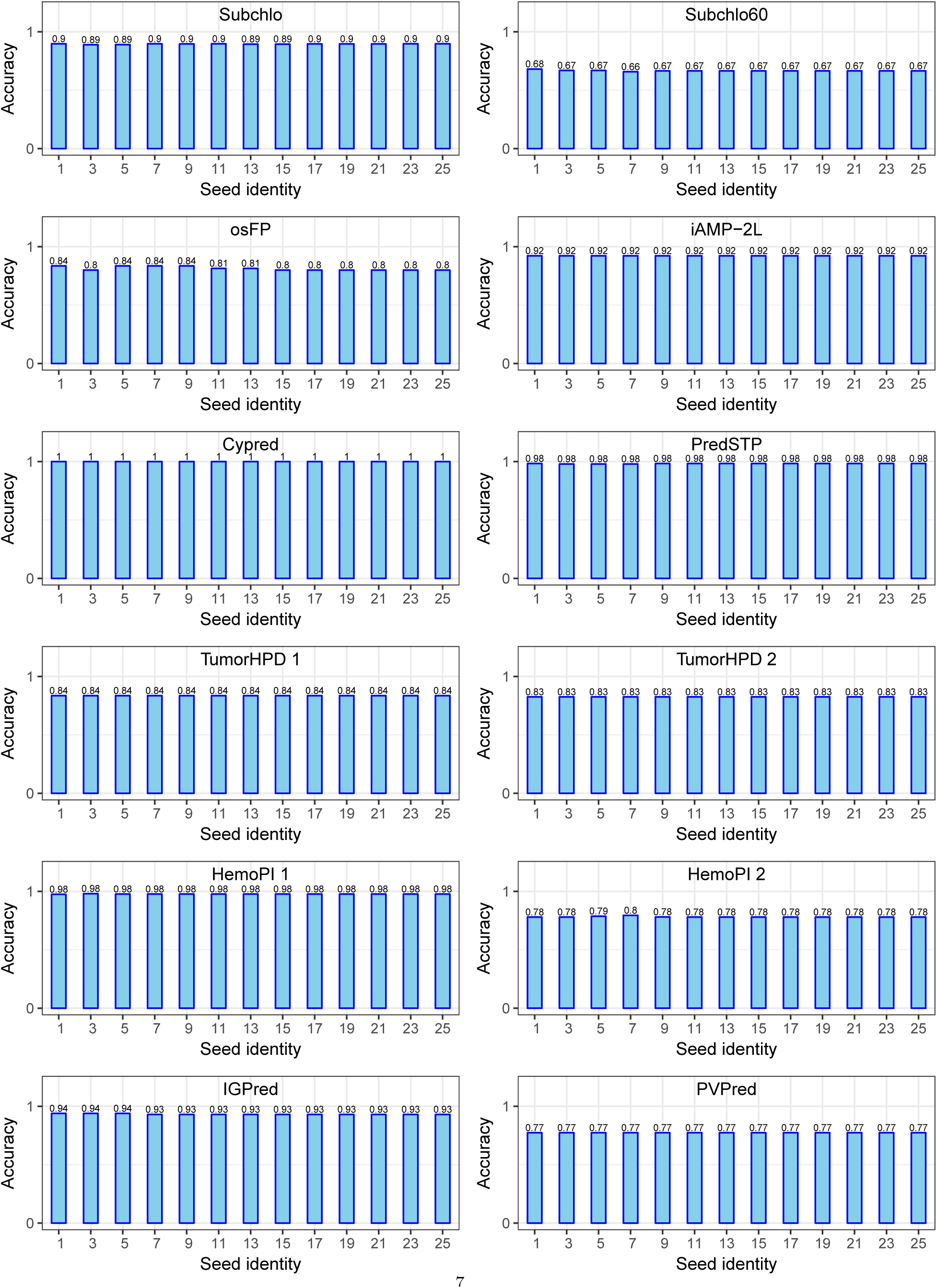
Represents the accuracies resulted from different seeds in a specific datasets. Each subplot represents an individual dataset. The x axis shows seed identities and the y axis shows the accuracy values.

**Fig. S3.**
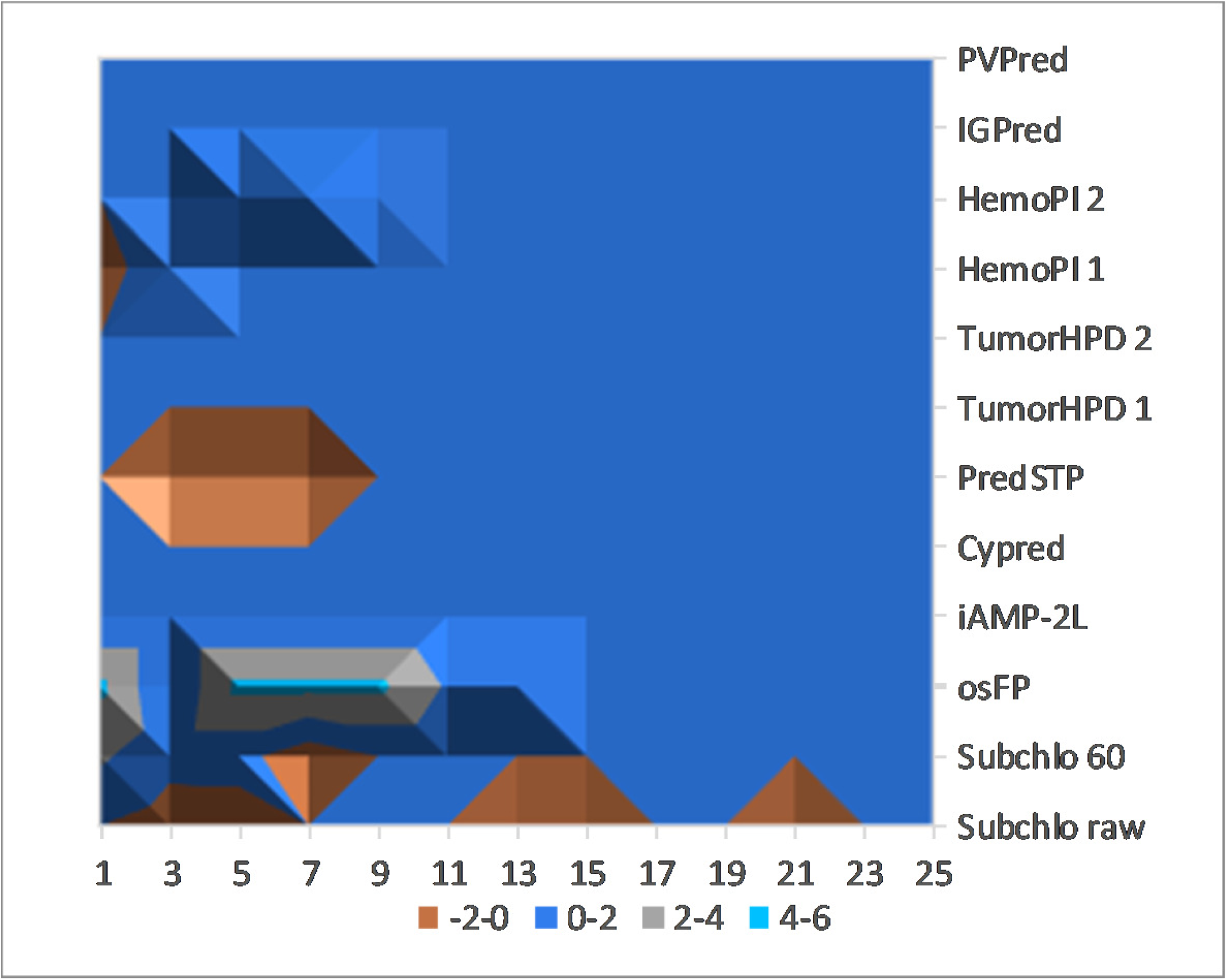
The percent change of accuracy for each seed compared to the mode accuracy of all the seeds for a specific dataset. The smaller the percentage deviation from the mode value, the better its convergence. iAMP-2L, Cypred, TumorHPD one, TumorHPD two, IGPred and PVPred showed perfect convergence, while the other datasets shows convergence for most of the seeds.

**Fig. S4.**
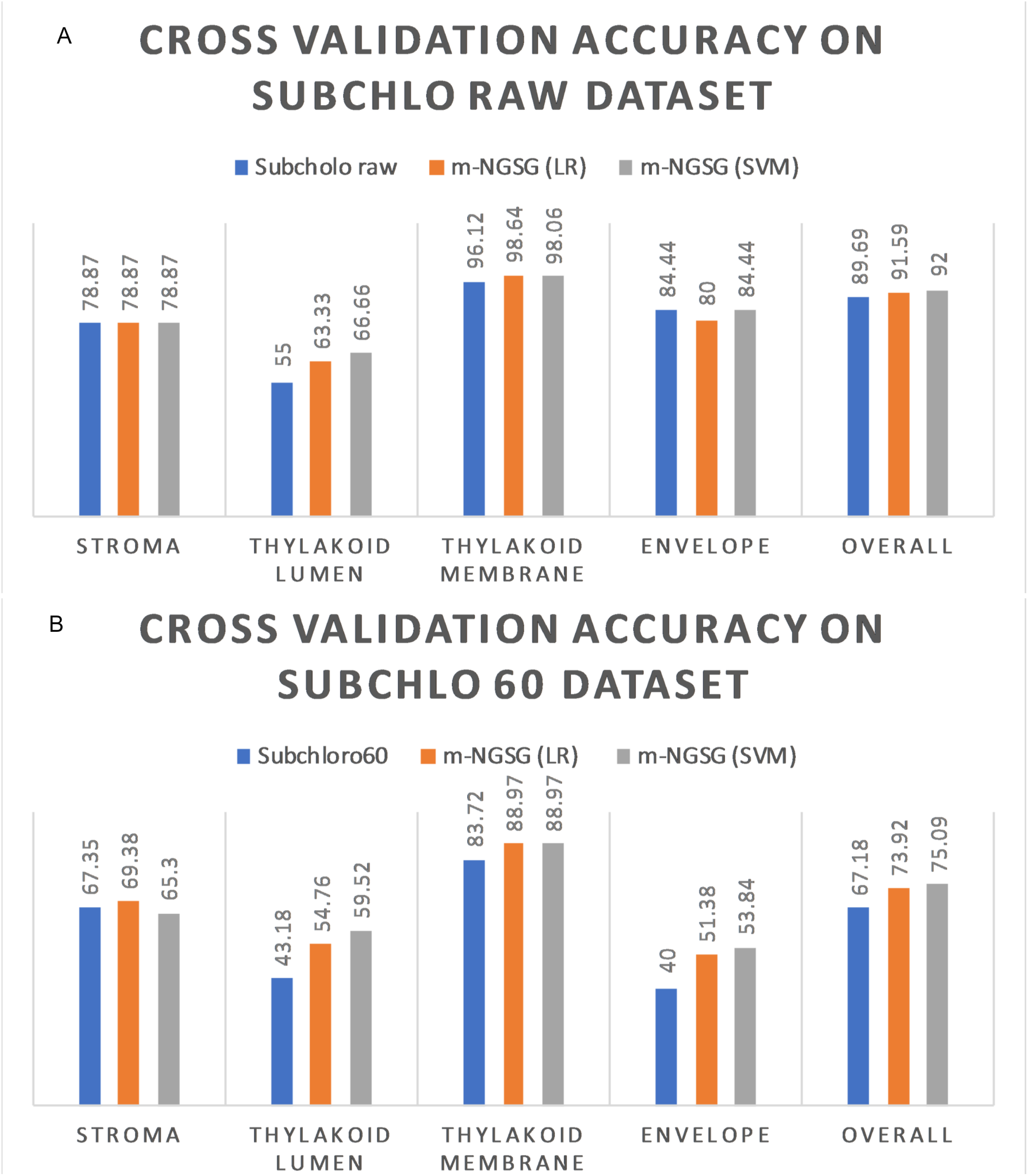
Cross-validation accuracy comparison between the original method and m-NGSG using Logistic Regression (LR) and linear kernel SVM (SVM) on Subchlo raw (A)and Subchlo60(B) datasets.

**Fig. S5.**
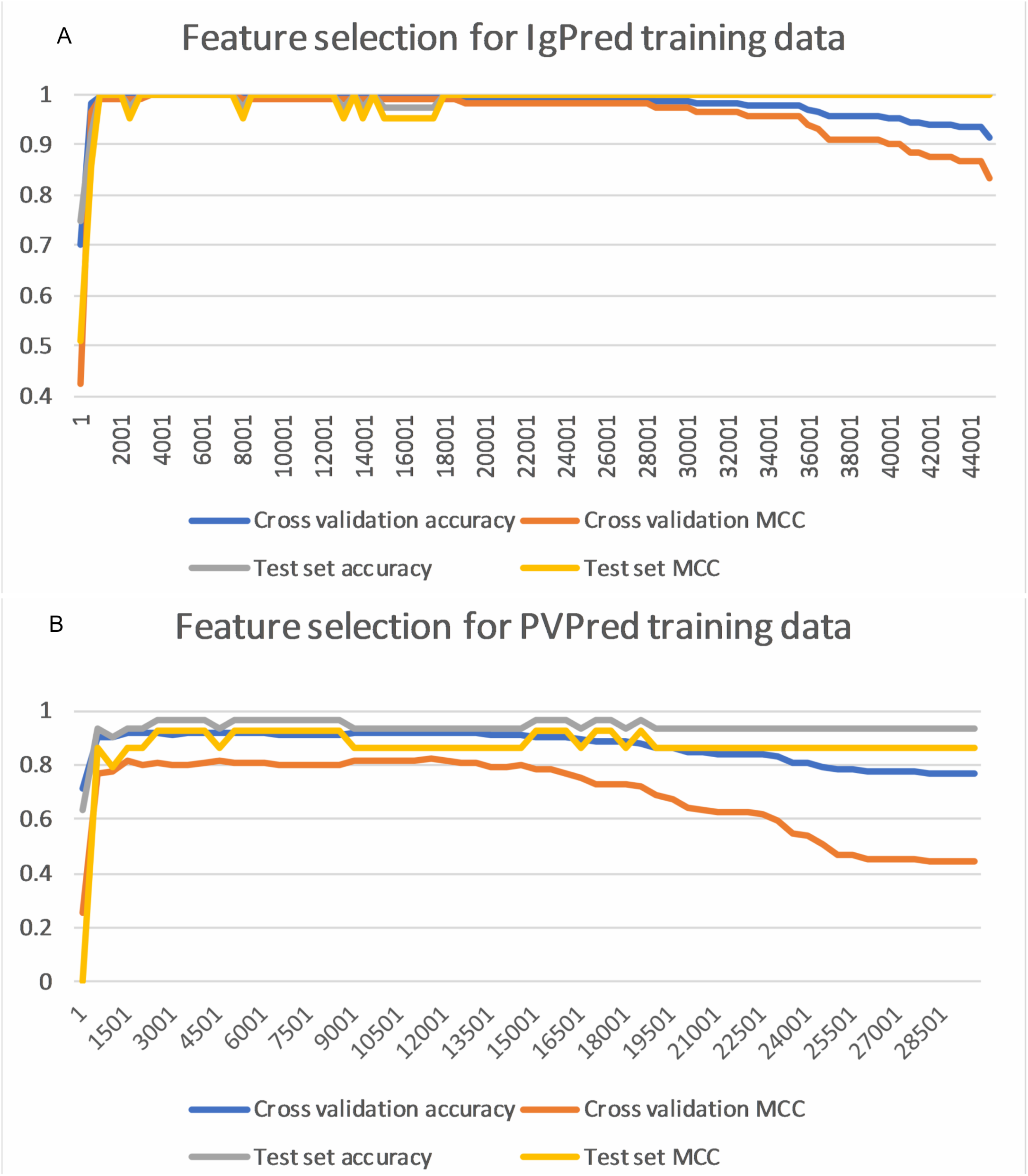
Performance of accuracies with number of features used. The features were added based on their importance according to ANOVA analysis. Using less features increase the cross-validation accuracy while a decrease of accuracy on the independent test set is evident as less features engender a bias cross validation. Figure A and B show the effect on accuracy and MCC values with increasing number of features on IGPred and PVPred datasets, respectively.

